# T cells induce interferon-responsive oligodendrocytes during white matter aging

**DOI:** 10.1101/2022.03.26.485917

**Authors:** Tuğberk Kaya, Nicola Mattugini, Lu Liu, Hao Ji, Simon Besson-Girard, Seiji Kaiji, Arthur Liesz, Ozgun Gokce, Mikael Simons

## Abstract

A hallmark of nervous system aging is a decline of white matter volume and function, but the underlying mechanisms leading to white matter pathology are unknown. Here, we found age-related alterations of oligodendrocytes with a reduction of total oligodendrocyte density in the aging murine white matter. Using single-cell RNA sequencing, we identify interferon-responsive oligodendrocytes, which localize in proximity of CD8^+^ T cells in the aging white matter. Absence of functional lymphocytes decreased oligodendrocyte reactivity and rescued oligodendrocyte loss, while T-cell checkpoint inhibition worsened the aging affect. In summary, we provide evidence that T cells induced interferon-responsive oligodendrocytes are important modifiers of white matter aging.

## MAIN TEXT

Age is the major risk factor for the most prevalent neurodegenerative diseases^1^. A better understanding of age-related alteration is therefore of importance, but relatively little is known about the pathology occurring in the white matter, which is to a large extent composed of myelin, a lipid-rich membrane wrapped around axons by oligodendrocytes^2^. Myelination is not limited to early development, but extends into adult life and contributes to brain plasticity. Regulated by neuronal stimuli and various environmental factors, there is a significant fraction of adult-born oligodendrocytes that are actively engaged in forming new myelin sheaths, a process that declines in aging^3,4,5^. In humans, white matter volume starts to decline already in mid-life, and these global alterations are often associated with focal lesions that appear hyperintense on magnetic resonance images^6,7^. Focal white matter degeneration is related to increased risk of stroke and dementia^6^, and contributes to cognitive decline possibly by disrupting connective pathways in the brain^8–10^. In non-human primates and in rodents, ultrastructural analyses of the aging white matter show pathology of myelinated fibers, consisting of focal areas of degenerated myelin^11,12^. We have previously shown that such age-related myelin pathology results in a distinct white-matter associated microglia, which activate partially the disease-associated microglia (DAM) or microglia-neurodegenerative phenotype (MGnD) program to clear myelin debris in groups of few closely connected microglia (nodules)^13–16^. The microglial responses to aging induced myelin pathology are intensely studied^13,15,17–19^, but much less is known about aging-related oligodendrocyte diversity that is producing myelin pathology. The aim of this study is to identify aging induced changes in oligodendrocytes.

To characterize oligodendrocyte aging, we performed single-cell RNA sequencing (scRNA-seq) to reveal transcriptomic cell-to-cell variation of oligodendrocytes in the normal and aged brain. We dissociated grey matter from the frontal cortex and white matter tracts from the corpus callosum as well as the optical tracts and the medial lemniscus, from young (3 months) and aged (24 months) old wild type mice (Fig. 1a). To avoid isolation artifacts, we used our previously established automated dissociation protocol that inhibits *ex-vivo* transcription^16^. We sorted live non-myeloid (CD11b^-^ and SYTOX^+^) cells for plate-based scRNA-seq (Smart-seq2), and analyzed after quality control the transcriptomes of 2538 single cells from 8 mice using unsupervised Uniform Manifold Approximation and Projection (UMAP) analysis (Fig. 1b; Supplementary Fig. 1c; Supplementary Tab. 1). Oligodendrocytes were separated in four different sub-clusters, of which the most abundant two clusters represent the heterogeneity of oligodendrocytes previously identified in juvenile and adult mouse by Marques et al^20^. In our analysis, two additional oligodendrocyte clusters appeared in aged mice, which were enriched in the white matter. One was characterized by a high expression of the serine (or cysteine) peptidase inhibitor, member 3N (Serpina3n), the complement component C4b, previously associated with injury responses^21–25^. Because this cluster was highly enriched in the aged white matter, we named it aging-related oligodendrocytes (Fig. 1b-d). We uncovered a smaller interferon-responsive oligodendrocyte subpopulation (IRO), which was characterized by the expression of genes commonly associated with a type I interferon response, such as *Stat1*, *Ifi27l2a* and major histocompatibility complex (MHC) class I-related genes (*H2-K1 and H2-D1*) (Fig. 1b, g-h). A related gene profile has been detected in oligodendrocytes progenitor cells (OPCs) in the context of multiple sclerosis^22,24^. To validate our results, we performed Drop-seq on 24 months old mice (8726 high-quality cells from 8 mice: Fig. 1b and Supplementary Fig. 1e). Tissues were prepared as described for Smart-seq2 and enriched for live cells using the flow-cytometry (Supplementary Fig. 1a). Major cell types were annotated based on canonical markers upon clustering (Supplementary Fig. 1e-g). We again identified a continuous range of oligodendrocytes that reproduced the major oligodendrocyte clusters of the Smart-seq2 scRNA-seq dataset (Fig. 1b, e). The higher number of oligodendrocytes allowed us to resolve Oligo1 and Oligo2 into five sub-clusters, but the identity and ratios of all four major clusters remained similar in both aged scRNA-seq datasets (Fig. 1d, f). The age-related oligodendrocytes and IRO were enriched in the white matter (Fig. 1f).

**Fig. 1.**
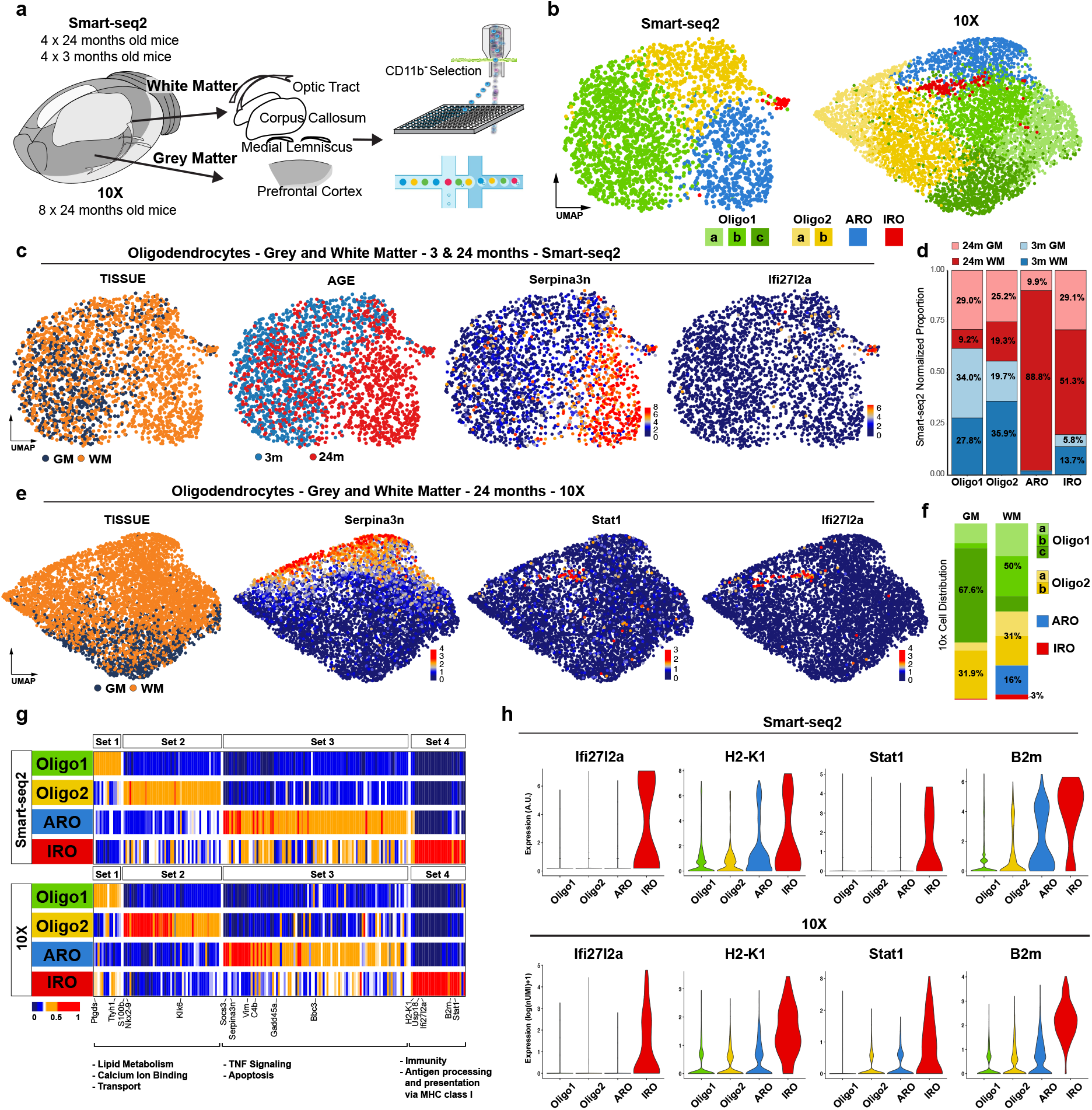
Identification of age-related gene expression signatures in oligodendrocytes. **a,** Experimental design from dissection to cell sorting and cell loading for the plate-based (Smart-seq2) and droplet-based (10X) pipelines, respectively. **b,** UMAP plots of oligodendrocytes in the SS2 and 10X datasets, colored by identified populations. **c,** UMAP plots of oligodendrocytes in the SS2 dataset, colored by tissue, age and expression of selected marker genes **d,** Bar plot showing the proportion of cells from each experimental group in each cluster. The number of cells from each experimental group was normalized to avoid biases due to total cell count discrepancies between groups. **e,** 10X dataset oligodendrocytes UMAP plots colored by tissue and expression of selected marker genes. **f,** Bar plot showing the relative distribution of each oligodendrocyte population in grey and white matter. **g,** Heatmaps of average expression of differentially expressed genes, comparing the 4 oligodendrocyte populations. Gene sets were identified for each population using the SS2 dataset. Values are normalized per gene, showing the gene expression across populations. Gene Ontology terms are shown below each set of genes. **h,** Violin plots showing selected IRO-enriched marker genes across SS2 and 10X datasets. A.U. is arbitrary units which represents the corrected log1p (counts) value.

To determine the localization and to validate the presence of the age-related and the interferon-responsive oligodendrocytes, we co-stained anti-adenomatous polyposis coli (APC) clone CC1 (CC1)^+^ oligodendrocytes by using antibodies against C4b, Serpina3n, STAT1 and B2M. Consistent with the scRNA-seq data, we found that antibodies against C4b, Serpina3n, STAT1 and B2M labeled oligodendrocytes in the white matter, and only rarely in the grey matter (cortical areas of the brain) of aged (24-month old) mice (Fig. 2a; Supplementary Fig. 2a, b). Co-labeling against STAT1 and Serpina3n did not detect double positive cells, in agreement with our scRNA-seq data that STAT1^+^ oligodendrocytes are distinct from Serpina3n^+^oligodendrocytes (Supplementary Fig. 3a, b). Next, we compared the labeling in young (3 months) and old (24 months) white matter tracts and found that CC1^+^ oligodendrocytes, also immunoreactive for C4b, Serpina3n, STAT1 or B2M, are restricted to the aged brains (Fig. 2a; Supplementary Fig. 2a, b). Quantification revealed that about 3 to 5% of the CC1^+^ cells within the corpus callosum of 24-month old mice were positive for the markers STAT1 and B2M. We found that C4b^+^/Serpina3n^+^ oligodendrocytes were not only more abundant (41% of CC1^+^ cells were Serpina3n^+^/CC1^+^ and 30% C4b^+^/CC1^+^in 24-month old mice white matter), but also more evenly spread as compared to B2M^+^/STAT1^+^ oligodendrocytes (Fig. 2a; Supplementary Fig. 4a). Our sub-regional localization analysis revealed that STAT1^+^ oligodendrocytes were often localized close to ventricular regions within the white matter (Supplementary Fig. 5d, e). Previous work has identified infiltrating T cells in the aged brain, close to neurogenic niches and within the optic nerve^26,27^. Because T-cells are one source of interferons, we analyzed the CD3^+^ T cells in the white and grey matter of the aged brain, and found that the T cells, which were mostly CD8^+^ T cells, were almost exclusively found in the white matter, where they were enriched in areas close to the lateral ventricles (Fig. 2b; Supplementary Fig.5d). Our previous work identified age-dependent formation of white-matter associated microglia, engaged in clearing myelin debris^16^. White-matter associated microglia are defined by the activation of genes implicated in phagocytic activity, lipid metabolism as well as antigen processing and presentation. First, we confirmed the age-dependent expression of major histocompatibility complex class I (MHC-I) in white-matter associated microglia nodules (Supplementary Fig. 6a-c). In addition, we found that microglia that localize to nodules were enriched in MBP transcripts (Supplementary Fig. 6d, e), reminiscent of microglia containing myelin transcripts previously detected in brains of multiple sclerosis patients^28^. Furthermore, microglial nodules were mostly in close proximity to CD8^+^ T-cells (Supplementary Fig. 6f-h). We have previously shown that triggering receptor expressed on myeloid cells 2 (TREM2) induces microglia nodule formation^16^. We therefore analyzed *Trem2*^-/-^ mice to determine possible differences in CD8^+^ T cell infiltration. For this analysis, we used 18 months wild type and *Trem2*^-/-^ mice, because this is the time point when CD8^+^ T cells start to appear. Indeed, the number of CD8^+^ T cells was markedly reduced in the corpus callosum of *Trem2*^-/-^ aged mice as compared to control, pointing to microglial involvement in CD8^+^ T cells infiltration (Supplementary Fig. 4d, e).

**Fig. 2.**
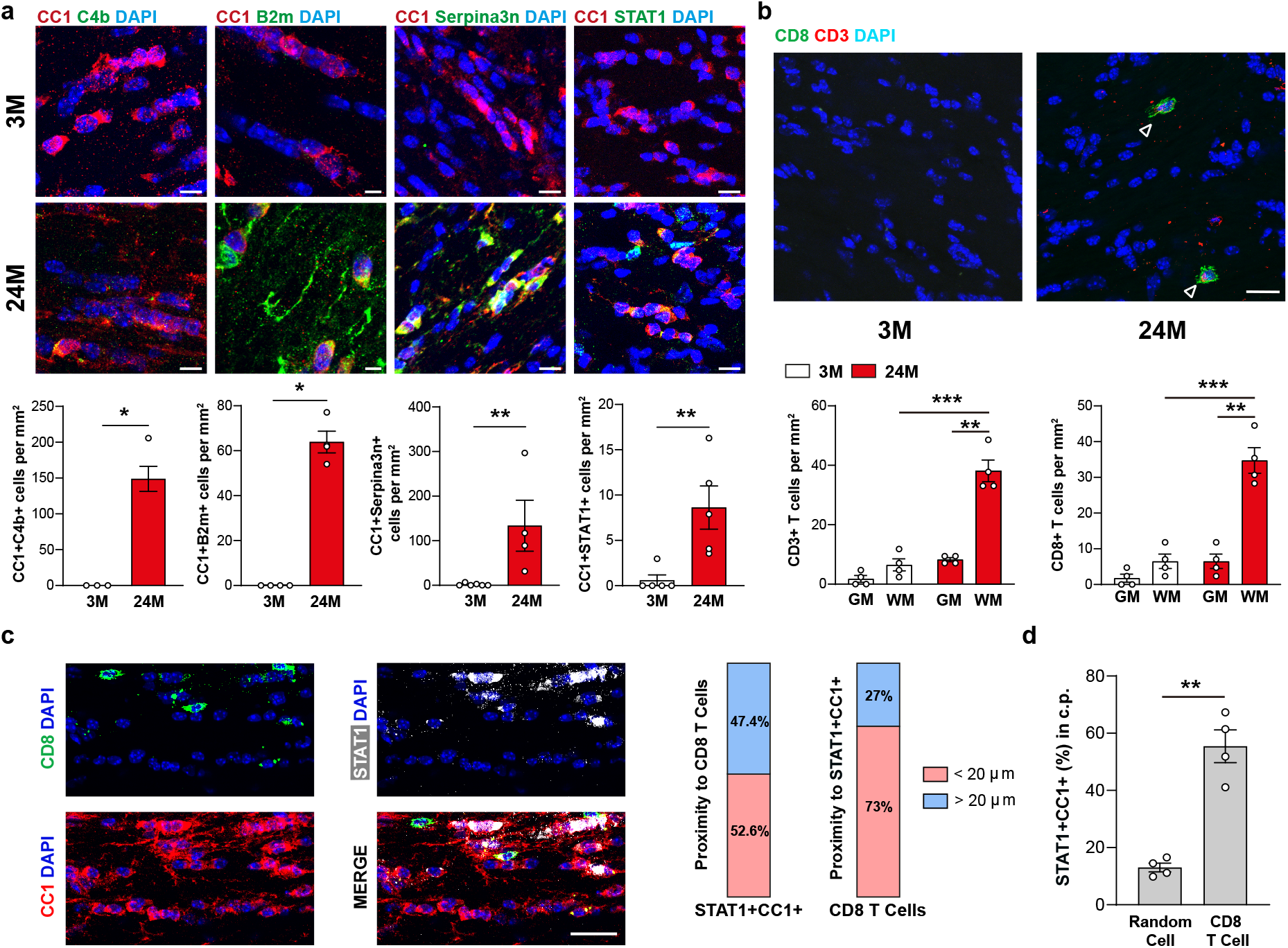
Interferon-responsive oligodendrocytes localize to the aged white matter close to CD8^+^ T-cells. **a,** Immunofluorescence staining showing expression of C4b, B2m, Serpina3n and STAT1 (green) in oligodendrocytes CC1^+^ (red) in the white matter of 3- and 24-month old mice. Scale bar for C4b, Serpina3n and STAT1, 20 μm, for B2m, 10 μm. Quantification of CC1^+^ oligodendrocytes expressing C4b, B2m, Serpina3n and STAT1 in the white matter of 3- and 24-month old mice (n=3-6 mice per group, mean ± sem; *P <0.05, **P < 0.01). **b,** Immunofluorescence showing T cells infiltration positive for CD3 (red) and CD8 (green) in the white matter. Arrows indicate CD3^+^CD8^+^ T cells of 3- and 24-month old mice. Scale bar, 20 μm. Quantification of CD3^+^ T cells and CD8^+^ T cells in the grey and white matter 3- and 24-month old mice (n=4 mice per group, mean ± sem; **P < 0.01, ***P < 0.001). **c,** Immunofluorescence of CD8^+^ T cells (green) and STAT1^+^CC1^+^ oligodendrocytes interaction in the white matter of 24-month old mice. Scale bars, 20 μm. The left bar plot shows the percentage of STAT1^+^CC1^+^ oligodendrocytes located within a 20 μm circle around the CD8^+^T cells (three sections per mouse were selected, a total of 134 CD8^+^ T cells from 4 mice were analyzed). The right bar plot shows the percentage of CD8^+^ T cells close to (<20 μm) STAT^+^CC1^+^ oligodendrocytes. **d,** Quantification of the percentage of STAT1^+^CC1^+^oligodendrocytes found around random cells (<20 μm) compared to the percentage of STAT1^+^CC1^+^ oligodendrocytes found around CD8^+^ T cells (three sections per mouse; n= 134 CD8^+^ T cells and 272 STAT1^+^CC1^+^ oligodendrocytes, n=134 random cells from 4 mice, mean ± sem.; *P < 0.05, **P < 0.01, ***P < 0.001).

Next, we studied the spatial relationship between the interferon-responsive oligodendrocytes (STAT1^+^CC1^+^) and CD8^+^ T cells in the aging white matter. Strikingly, STAT1^+^CC1^+^ and CD8^+^T cells are frequently localized in close proximity (less than 20 μm) (Fig. 2c). Moreover, STAT1^+^CC1^+^ oligodendrocytes are significantly more often found in close proximity to CD8^+^T cells than randomly chosen DAPI^+^ cells (Fig. 2d).

To determine the functional impact of CD8^+^ T cells on oligodendrocytes, we treated 18-month old mice with antibodies against the co-inhibitory receptors such as Cytotoxic T-Lymphocyte-Associated Protein 4 (CTLA-4) and Programmed cell death protein 1 (PD-1). Immune checkpoint blockage therapy is used to dampen co-inhibitory molecules to achieve robust antitumor immune response^29^. Treatment of mice with two times a week i.p. injections with anti–PD-1 and anti–CTLA-4 antibody for six weeks resulted in an increase of CD8^+^ T cells in the white matter (Fig. 3a, b). In addition, the number of STAT1^+^CC1^+^ and B2M^+^CC1^+^oligodendrocytes increased within the corpus callosum by checkpoint blockage therapy (Fig. 3b). Notably, the number of Serpina3n^+^ oligodendrocytes was unaffected by anti–PD-1/anti–CTLA-4 antibody treatment (Supplementary Fig. 4b, c).

**Fig. 3.**
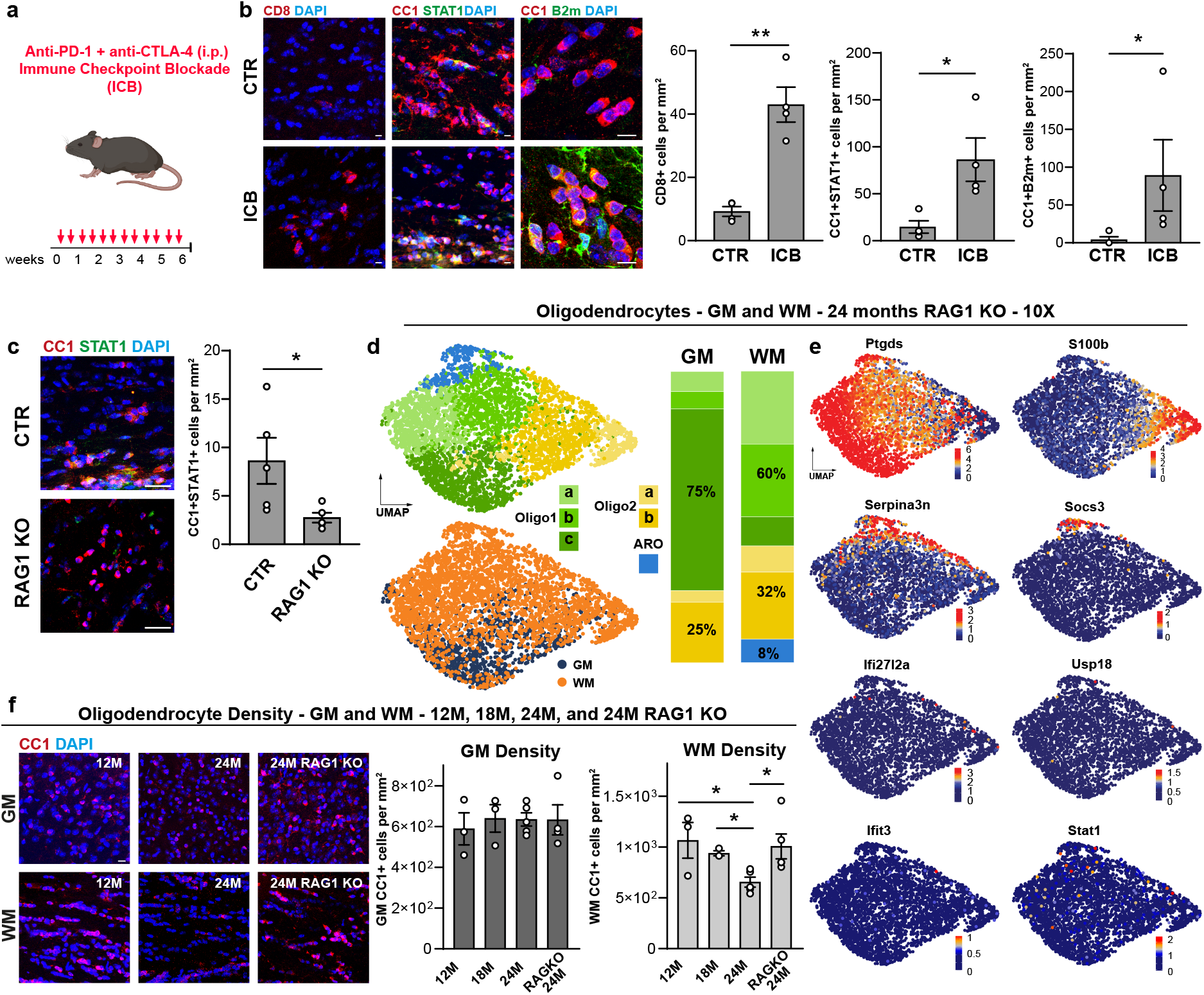
Absence of functional lymphocytes reduces interferon-responsive oligodendrocyte number and increases oligodendrocyte cell density in the aged white matter. **a,** Graphical presentation of experimental outline. **b,** Immunofluorescence showing CD8^+^ T cells in the white matter of mice treated with anti PD-1 and CTLA-4 and isotype control antibodies for 6 weeks starting at an age of 18 months. Immunofluorescence showing B2m and STAT1 (green) in CC1^+^ oligodendrocytes (red) in the white matter of mice treated with antibodies anti PD-1 and CTLA-4 and isotype control antibodies. Scale bar, 20 μm. Quantification CD8^+^ T cells in the white matter of mice treated with antibodies anti PD-1 and CTLA-4 and isotype control antibodies (n=4 mice per group, mean ± sem; *P < 0.05, **P < 0.01). Quantification of CC1^+^ oligodendrocytes expressing B2m and STAT1 in the white matter of mice treated with antibodies anti PD-1 and CTLA-4 and isotype control antibodies (n=4 mice per group, mean ± sem.; *P < 0.05). **c,**Immunofluorescence showing expression of STAT1 (green) in oligodendrocytes CC1^+^ (red) in the white matter of mice 24-month old wildtype and 24-month old *Rag1*^-/-^ mice. Scale bar, 20 μm. Quantification of CC1^+^STAT1^+^ oligodendrocytes in the white matter of mice 24-months old wild type and 24-month old *Rag1*^-/-^ mice (n=5 mice per group, mean ± sem; *P < 0.05). **d,** UMAP plots of oligodendrocytes in the *Rag1*^-/-^ dataset. Identified clusters and tissue annotation are color coded, in the upper and lower plots, respectively. Bar plot showing the relative distribution of each oligodendrocyte population in grey and white matter. **e,** UMAP plots of the expression of selected marker genes. **f,** Immunofluorescence showing CC1^+^ oligodendrocytes (red) in the grey and white matter of 12- and 24-month old wild type and *Rag1*^-/-^ mice. Scale bar, 20 μm. Quantification of CC1^+^oligodendrocytes in the grey and white matter of 12-, 18-, 24-month old wild type and *Rag1*^-/-^ mice (n=3-5 mice per group, mean ± sem; *P < 0.05).

To continue exploring the link between CD8^+^ T cells and STAT1^+^CC1^+^ oligodendrocytes, we investigated aged *Rag1*^-/-^ mice. We found that the absence of functional lymphocytes decreased the number STAT1^+^CC1^+^ oligodendrocytes in 24-months old *Rag1*^-/-^ mice as compared to control wild-type mice (Fig. 3c). Next, we subjected aged white and grey matter of *Rag1*^-/-^ mice to scRNA-seq to determine systemically how oligodendrocyte subpopulations were altered in the absence of lymphocytes. Unsupervised clustering of combined datasets showed that oligodendrocytes distributed in different clusters with the same transcriptional profile that we had identified in the wild-type datasets, excluding the IRO population (Fig. 3d, e). This finding was consistent with our immunolabeling experiments, which showed a marked reduction of IROs in *Rag1*^-/-^ mice (Fig. 3c).

Together, these data provide evidence that the adaptive immune system promotes oligodendrocytes reactivity in the aging white matter, but to what extent these changes contribute to white matter degeneration is unclear. First, we determined whether the density of oligodendrocytes changes in the aging brain. Because oligodendrocytes are continuously generated to an age of around 12 months in mice, we quantified oligodendrocytes cell number in 12-, 18- and 24-month old mice (Fig. 3f). We found that the density of CC1^+^ oligodendrocytes declined in the white matter, whereas no changes were observed in the aged grey matter (Fig. 3f). Similar results were obtained when Glutatione S-Transferase (GST-π) was used as an additional marker to stain for mature oligodendrocytes (Supplementary Fig. 7a, b). Quantification showed that both CC1^+^ and GST-π^+^ cells decreased in the 24-month old corpus callosum compared to 12-month, while the number remained stable in the grey matter (Fig. 3f; Supplementary Fig. 7a, b). In addition, there was a decrease in OLIG2^+^ cells of the oligodendrocyte lineage (Supplementary Fig. 7a, b). Next, we determined the density of oligodendrocytes in the aged white matter of *Rag1*^-/-^ mice. There was a higher density of oligodendrocytes in the aged white matter in *Rag1*^-/-^ compared control mice (Fig. 3c).

In summary, we provide evidence that T cells contribute to the cellular alterations that are associated with white matter aging. White matter aging causes myelin degeneration, which results in microglia reactivity with a fraction of cells accumulating in nodules where they are engaged in myelin debris clearance. We now find that aging is also associated with oligodendrocyte responses, as seen by the generation of a subpopulation of STAT1^+^ and Serpina3n^+^/C4b^+^ oligodendrocytes in the aging white matter. In addition, we find a reduction of oligodendrocyte density in the aged white but not grey matter. Absence of functional lymphocytes decreases the number STAT1^+^/B2M^+^ oligodendrocytes and increases the total density of oligodendrocytes, providing evidence that the adaptive immune system is an important modifier of white matter aging. The temporal sequence of pathological processes with the aging white matter still needs to be established. As the deep white matter areas lie at the ends of the arterial circulation, they are particularly susceptible to decreases in blood flow and oxygenation, possibly contributing to increased vulnerability of aged white matter to hypoperfusion and aging induced leaky blood brain barrier^30,31^. Progressive vascular damage to myelinated fibers may not only result in microglia reactivity, required to clear the increasing amounts of myelin debris that accumulate during aging, but also to infiltration of CD8^+^ T cells, thereby triggering harmful immune reactivity towards oligodendrocytes. Previous work has shown that oligodendrocytes are particularly sensitive to interferon-γ, as it is able to trigger oligodendroglial cell death and demyelination^32,33^. Immune responses in the aging brain are not limited to oligodendrocytes. Previous work has shown T-cell infiltration in the aged brain, where they impair the function of neuronal stem cells within the neurogenic niche and induce damage to axons in the optic nerve^26,27,34^. The clonal expansion of CD8^+^ T cells provides evidence that they actively recognize antigen(s), and may not only simply migrating into the brain because of age-related disruption of the blood-brain barrier^27^. An open future question is whether and if so, which antigens CD8^+^ T cells recognize in the aged white matter. Proof-of-principle that oligodendrocyte pathology is able to trigger adaptive autoimmune responses against myelin has been provided in a model of oligodendrocyte ablation^35^. Even if the exact link between CD8^+^ T cells and oligodendrocytes remains to be established, these results raise the intriguing possibility that cytotoxic CD8^+^ T cells contribute to decline of white matter function in aging.

## SUPPLEMENTARY FIGURES

**Supplementary Figure 1.**
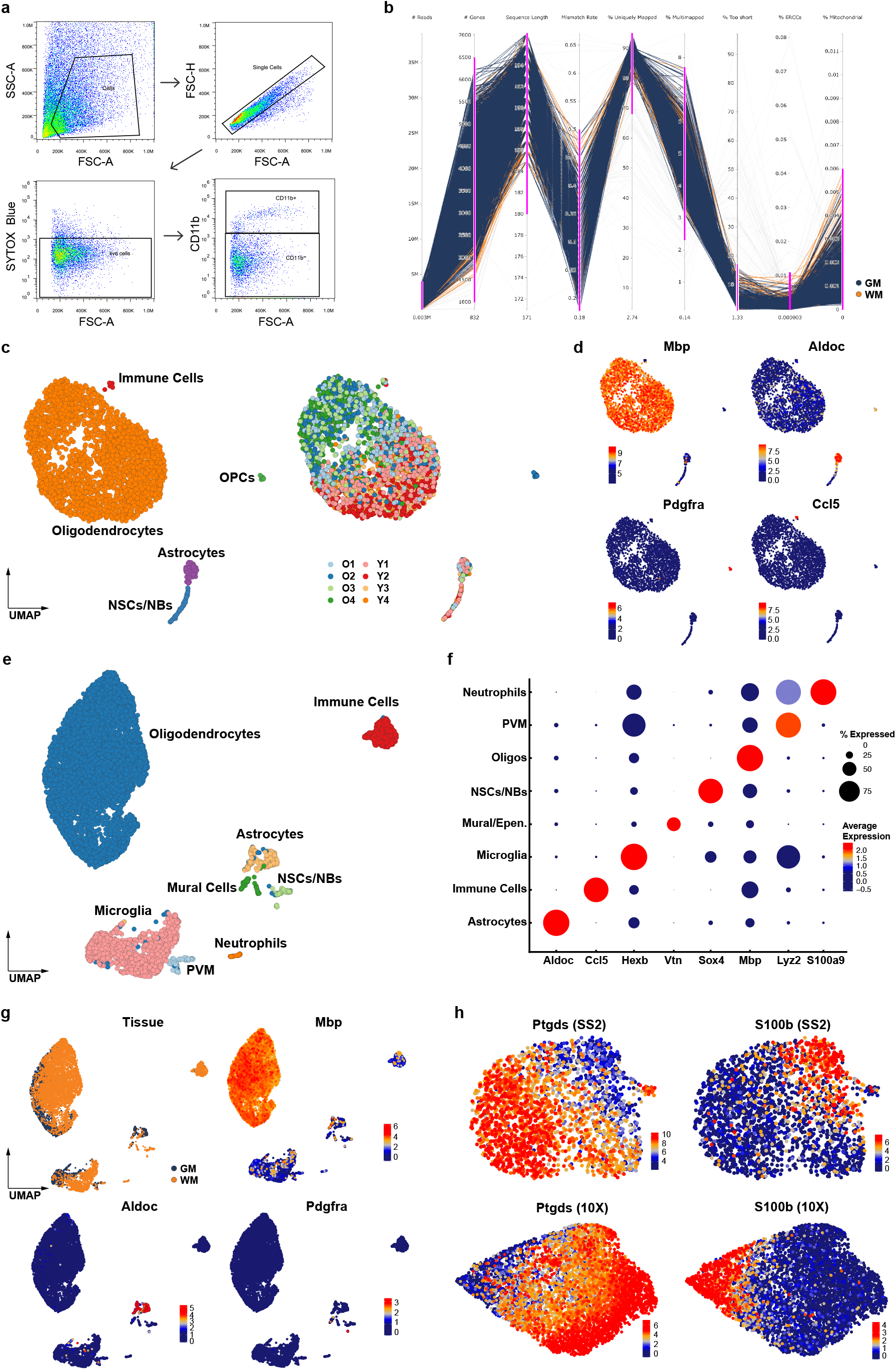
**a,** Sorting strategy for the SS2 libraries. Flow cytometry gating of CD11b negative cells to enrich for oligodendrocytes and astrocytes. **b,** Quality control (QC) for the SS2 dataset. Quantitative criteria are represented along with the chosen thresholds (coloured in pink for each step). Out of 2650 sequenced single-cells, 2538 passed the set of quantitative QC metrics. **c,** UMAP plots of 2538 single-cell transcriptomes. Cell type clusters and samples from different animals are colour coded. O1-2-3-4 and Y1-2-3-4 correspond to samples from 24-month old and 3-month old animals, respectively. **d,** UMAP plots of selected cell typespecific marker genes in the SS2 dataset; *Mbp* (Oligodendrocyte), *Aldoc* (Astrocyte), *Pdgfra* (OPC), *Ccl5* (Immune cells). **e,** UMAP plot of 8726 single-cell transcriptomes in the 10X dataset. Different cell type clusters are color coded. **f,** Dot plot showing cell-type marker gene expression by clusters. **g,** UMAP plots of tissue annotation and cell-type specific marker genes in the 10X dataset. **h,** UMAP plots of the oligodendrocyte populations depicting the expression of Oligo1 and Oligo2 marker genes (*Ptgds* and *S100b*, respectively), in the SS2 and 10X dataset.

**Supplementary Figure 2.**
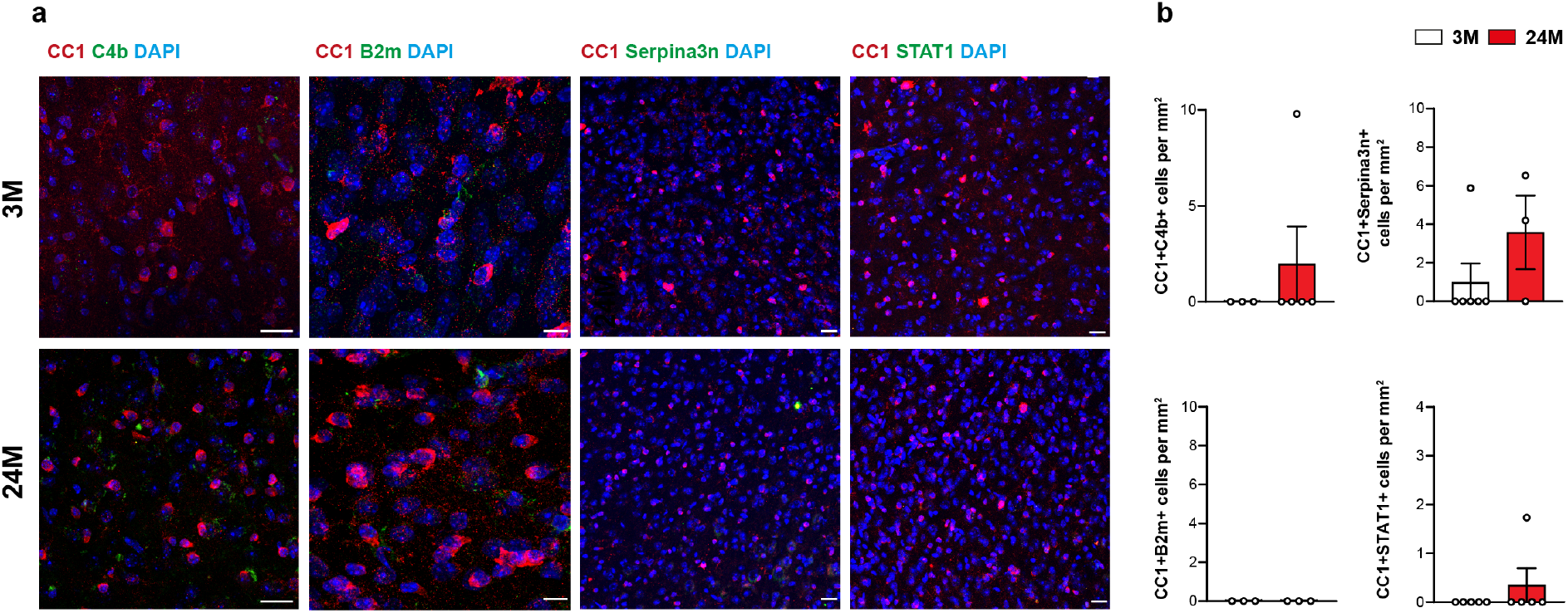
**a,** Immunofluorescence showing expression of C4b, B2m, Serpina3n and STAT1 (green) in CC1^+^ oligodendrocytes (red) in the grey matter of 3- and 24-month old mice. Scale bar for C4b, Serpina3n and STAT1 20 μm, for B2m 10 μm. **b,** quantification of CC1^+^ oligodendrocytes expressing C4b, B2m, Serpina3n and STAT1 at 3- and 24-month old mice in the white matter (n=3-6 mice per group, mean ± sem)

**Supplementary Figure 3.**
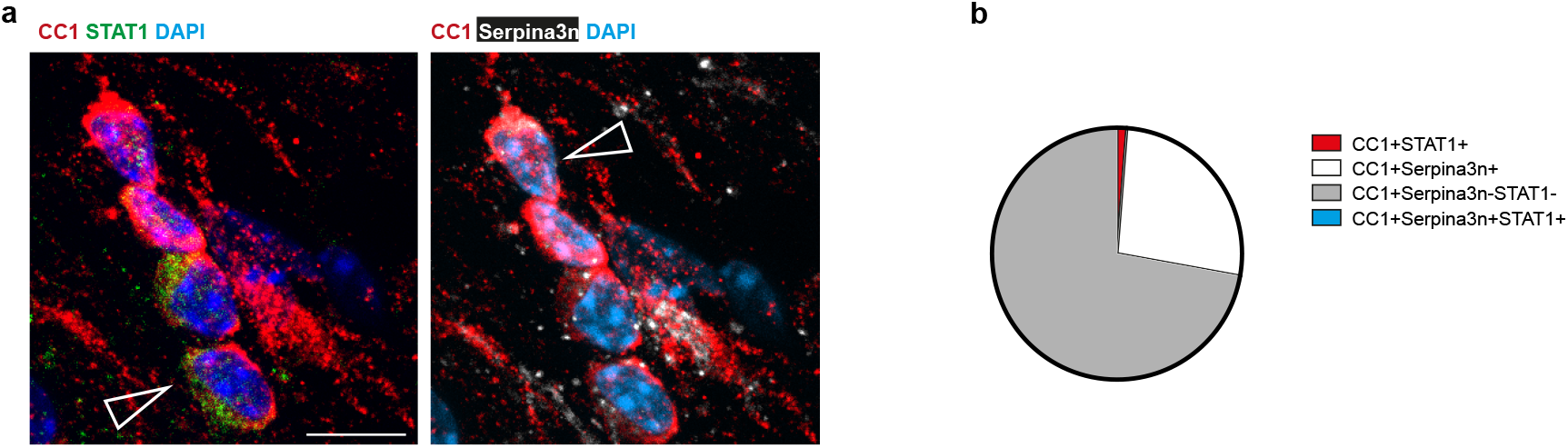
**a,** Immunofluorescence showing expression Serpina3n (white) and STAT1 (green) in CC1^+^ oligodendrocytes (red) in 24-month old mice in the white matter. Scale bar, 10 μm. Arrows indicate double cells for STAT1^+^CC1^+^ and Serpina3n^+^CC1^+^. **b,** Pie chart of percentage of different subpopulations.

**Supplementary Figure 4.**
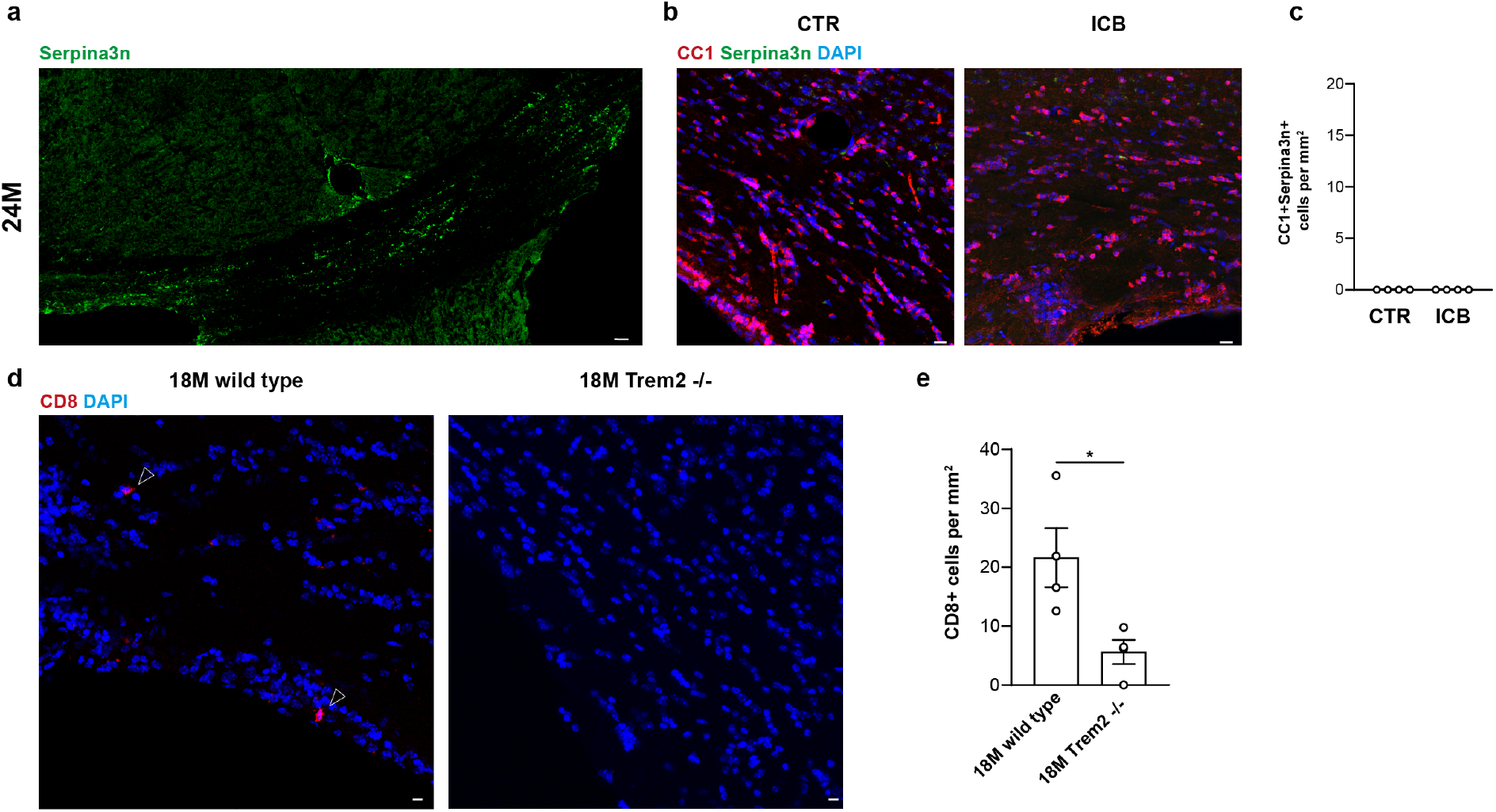
**a,** Immunofluorescence of Serpina3n distribution in the white matter of 24-month old mice. Scale bar, 50 μm. **b,** Immunofluorescence showing expression of Serpina3n (green) in CC1^+^ oligodendrocytes (red) in the white matter of mice treated with antibodies anti PD-1 and CTLA-4 and isotype control for 6 weeks starting at an age of 18 months. Scale bar 20 μm. **c,** Quantification of CC1^+^ oligodendrocytes expressing Serpina3n in the white matter of mice treated with antibodies anti PD-1 and CTLA-4 and isotype control antibodies (n=4 mice per group, mean ± sem). **d,** Immunofluorescence of CD8^+^ T (red) cells in the white matter of 18-month old wild type and *Trem2*^-/-^. Arrows indicate CD8^+^ T cells. Scale bar 20 μm. **e,** Quantification of CD8^+^ T cells in the white matter of 18-month old wild type and *Trem2*^-/-^ (n=4 mice per group, mean ± sem; *P < 0.05).

**Supplementary Figure 5.**
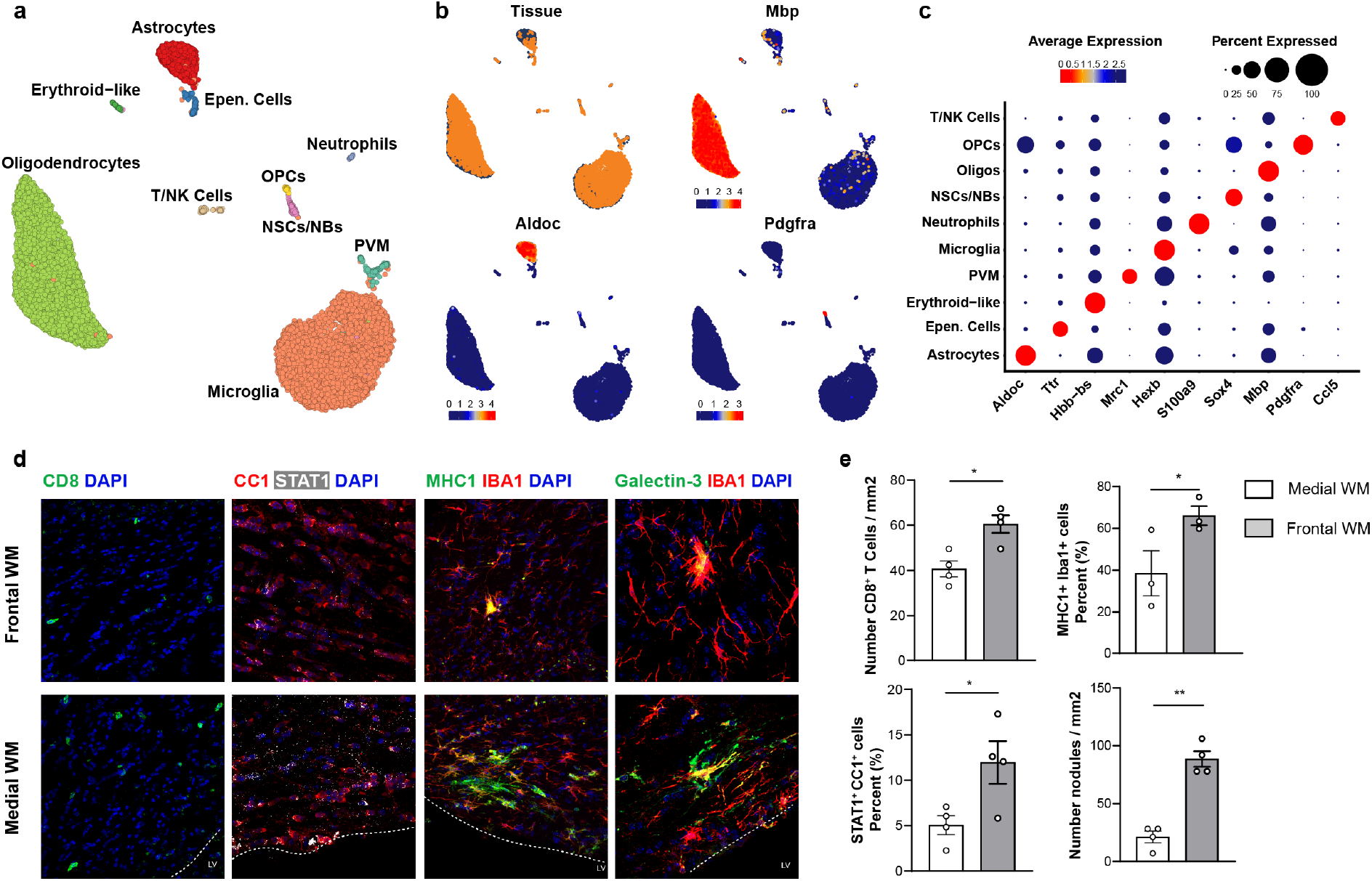
**a,** UMAP plot of 11438 single-cell transcriptomes in the *Rag1*^-/-^ dataset. Different cell type clusters are color-coded. **b,** UMAP plots of tissue annotation and cell-type specific marker genes in the *Rag1*^-/-^ dataset. **c,** Dot plot showing cell-type marker gene expression by clusters. **d,** Immunofluorescence staining showing the different densities of CD8^+^T cells, STAT1^+^CC1^+^ cells, MHC1^+^IBA1^+^ cells and nodules (Galectin-3^+^Iba1^+^ cell clusters) between frontal white matter and medial white matter. Scale bar 20 μm. **e,** Quantification of CD8^+^ T cells, STAT1^+^CC1^+^ cells, MHC1^+^IBA1^+^ cells, STAT1^+^CC1^+^ cells and nodules (Galectin-3^+^IBA1^+^ cells cluster) density difference between frontal white matter and medial white matter of 24-month old mice (3-4 mice, two section per group from each mice, mean ± sem; *P < 0.05, **P < 0.01).

**Supplementary Figure 6.**
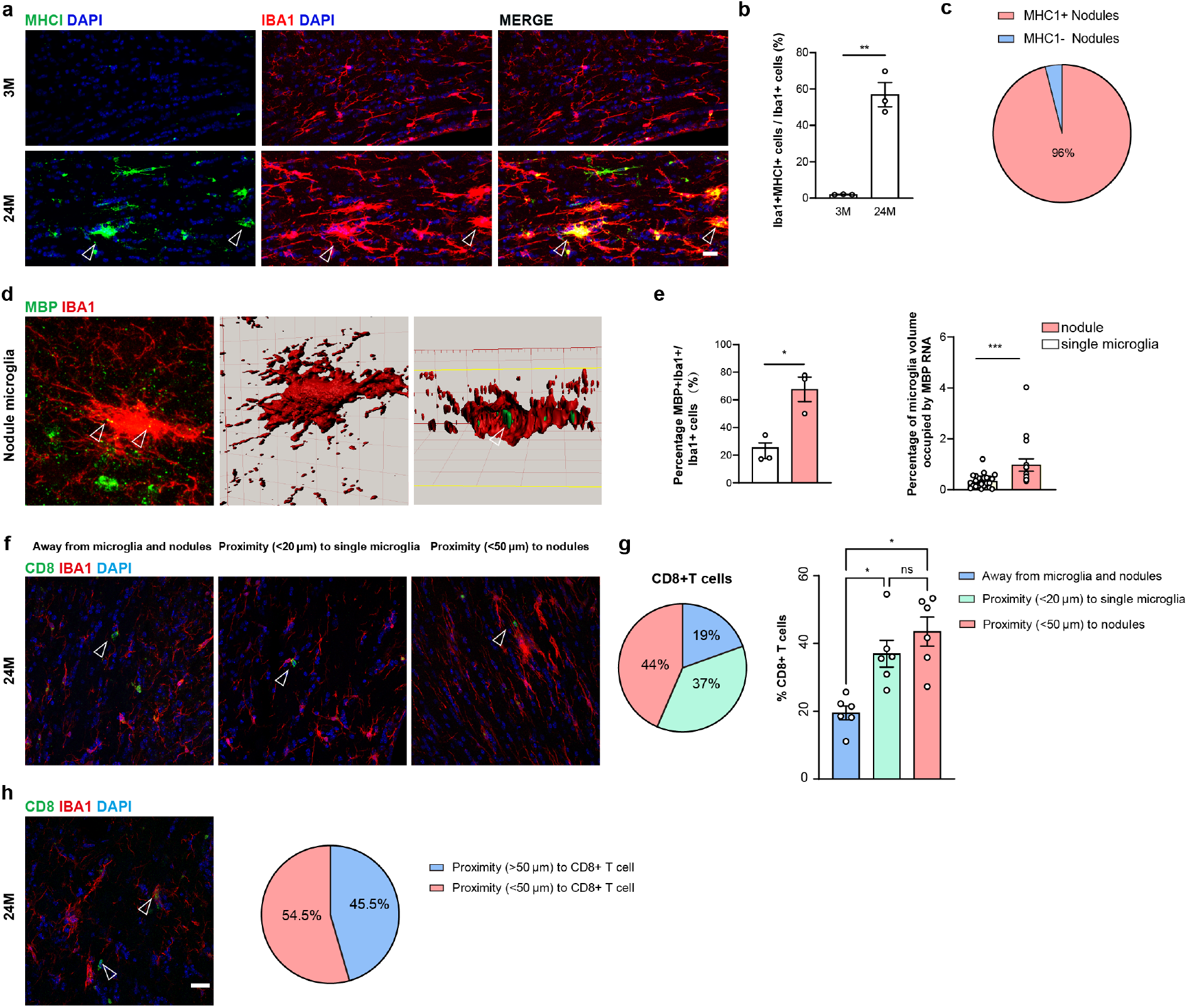
**a,** Immunofluorescence showing expression MHC1 (green) in microglia IBA1^+^ (red) in the white matter of 3-month old and 24-month old mice. Arrow marks MHC1^+^ nodules. Scale bar, 20 μm. **b,** Quantification of IBA1^+^ microglia expressing MHC1 in the white matter of 3- and 24-month old mice (n=3 mice per group, mean ± sem; **P <0.01). Arrows mark double positive microglia. **c,** Pie chart showing that most of the nodules are MHC1^+^ (60 IBA1^+^ nodules from 3 mice were analyzed**). d,** Immunofluorescence combined with RNAscope of MBP mRNA (green) within IBA1^+^ microglia nodule (red). Clipped 3D images show MBP mRNA in a microglia nodule from a different angle. Arrow mark MBP mRNA inside microglia. Scale bar, 20 μm, 3D rendering 10 μm. **e,** Quantification of percentage of IBA1^+^ single microglia and IBA1^+^ nodule with MBP mRNA inside (n=3 mice per group, mean ± sem; *P < 0.05). Quantification of percentage of IBA1^+^ volume occupied by MBP mRNA (single microglia = 26 cells, 30 IBA1^+^ nodules; ***P < 0.001). **f,** Immunofluorescence staining showing the three different spatial position relations of CD8^+^ T cells to microglia in the white matter of 24-month old mice: T cells away from microglia and nodules, T cells proximity (<20 μm) to single microglia, and T cells in close proximity (<50 μm) to nodules. Scale bar, 20 μm. **g**, Pie chart of the percentage of CD8^+^ T cells in different spatial groups as seen in panel f. Quantification of the percentage of CD8^+^ T cells distribution between three groups (337 CD8^+^ T cells from 6 mice were analyzed, similar number of T cells from each mouse were selected; mean ± sem; *P < 0.05). **h,** Immunofluorescence and a pie chart showing that more than half of CD8^+^ T cells (green) are in proximity (<50 μm) to nodules (IBA1^+^, red). Arrows mark CD8^+^ T cells. A total number of 122 IBA1^+^ nodules from 6 mice were analyzed. Scale bar, 20 μm.

**Supplementary Figure 7.**
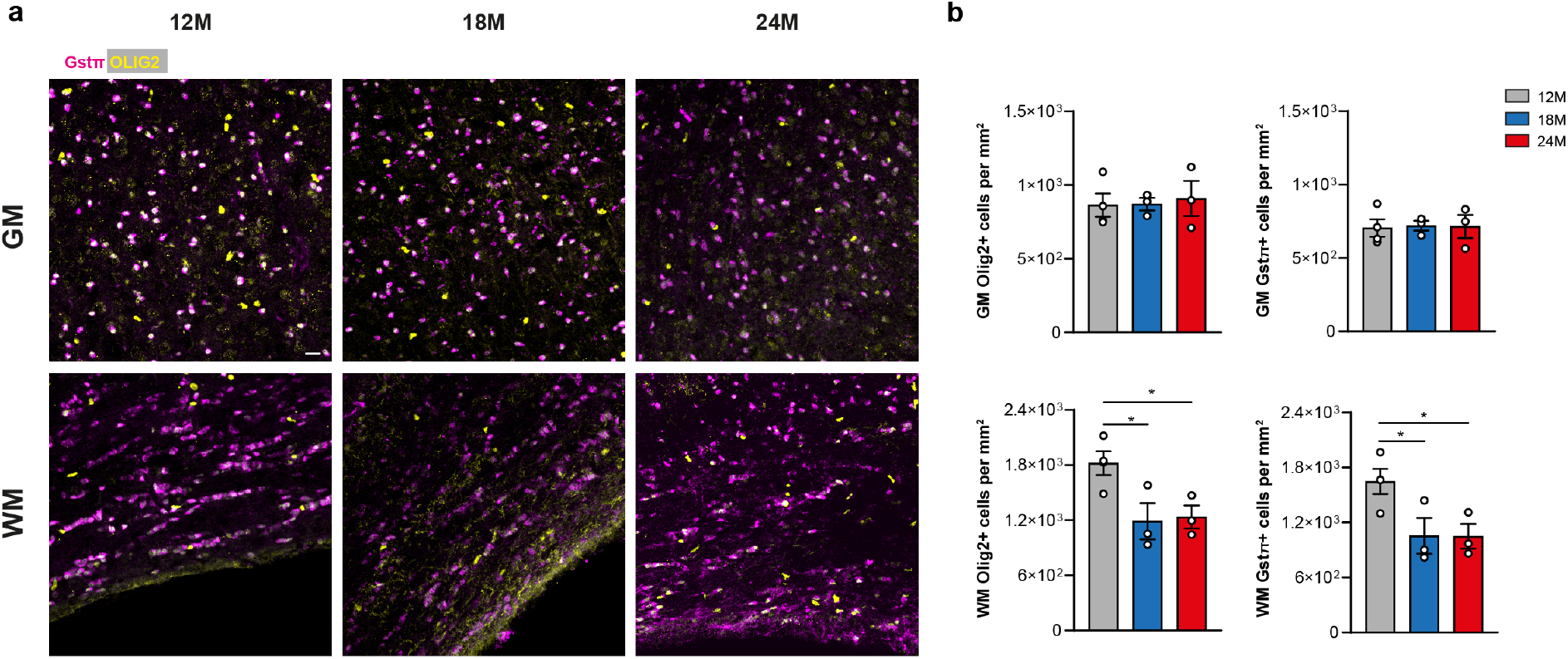
**a,** Immunofluorescence showing OLIG2^+^ oligodendrocytes lineage cells (yellow) and oligodendrocytes GSTπ^+^ (magenta) and in the grey and white matter of 12-, 18-, 24-month old mice. Scale bar 20 μm. **b,** Quantification OLIG2^+^ and GSTπ^+^ cells in the grey and white matter of 12-, 18-, 24-month old mice (n=3-4 mice per group, mean ± sem; *P < 0.05).

**Supplementary Table 1|** A complete list of mice used for scRNA-seq.

## ACKNOWLEDGMENTS

The work was supported by grants from the German Research Foundation (SPP2191, TRR 128-2, Project ID 408885537-TRR 274, SyNergy Excellence Cluster, EXC2145, Projekt ID390857198), the Human Frontier Science Program (HFSP), the ERC (Consolidator Grant), and the Dr. Miriam and Sheldon G. Adelson Medical Research Foundation, Chan-Zuckerberg Initiative grant, and Else Kröner-Fresenius-Stiftung grant. For the single cell and sorting studies, we are grateful for support from the “Flow Cytometric Cell Sorter Sony SH800 Core Unit” run by the Department of Vascular Biology at the Institute for Stroke and Dementia Research (ISD) and SyNergy EXC2145. We thank Christian Haass for providing TREM2 knockout mice.

## AUTHOR CONTRIBUTIONS

M.S., O.G. conceived and supervised the project. T.K., N.M., L.L., S.B.-G., H.J., S.K., O.G. performed experiments and analyzed the data, M.S., O.G. analyzed the data or supervised data acquisition, A.L. provided essential reagents, M.S., O.G. wrote the manuscript with input from all authors.

## METHODS

### Animals

The mouse line used in this study are the following: Wild-type C57BL/6J mice were from Janvier Labs; *Trem2*^-/-^ mice^36^ on the C57BL/6J background were kindly provided by Prof. Christian Haass, Laboratory of Neurodegenerative Disease Research, DZNE, Munich; *Rag1*^-/-^ were on the C57BL/6J background and purchased from the Jackson Laboratory. Experiments were performed with adult mice of 3, 12, 18 and 24 months as indicated in the figures. Treatment with antibodies against PD-1 and CTLA-4 and their respectively isotype control were performed on adult mice of 18-month old; mice were injected intraperitoneally (i.p.) with a mix of both antibodies with a concentration of 10 mg/Kg (PD-1) and 20 mg/Kg (CTLA-4) twice a week for 6 weeks in total. All animal experiments were reviewed and overseen by the institutional animal use and care committee in German Center for Neurodegenerative Diseases (DZNE) in Munich. The mice were kept in groups in Greenline IVC GM500 plastic cages and were housed in a temperature-controlled environment (21 ± 2C) on a 12 h light/dark cycle with food and water available ad libitum in the animal facility in German Center for Neurodegenerative Diseases (DZNE) in Munich.

### Mice perfusion, cell isolation for Smart-seq2

Four young (3-month old), and four old (24-month old) male C57BL/6 mice were deeply anesthetized and perfused with cold PBS (Sigma, D8537). Each brain was carefully removed and individually micro-dissected under a dissection microscope; grey matter was isolated from the frontal cortex and white matter was carefully isolated from the optic tract, medial lemniscus and corpus callosum (attached grey matter and choroid plexus were removed). We used our previous established isolation protocol^37^ using gentleMACS™ with the Neural Tissue Dissociation Kit (Papain) (Miltenyi Biotec) and a final concentration of 45 mM actinomycin D (Act-D, Sigma, No.A1410). Subsequently, cells were blocked with mouse FcR-blocking reagent (CD16/CD32 Monoclonal Antibody, eBioscience cat:14-0161-82,1100), stained with antibodies against CD11b (PE/Cy7,M1/70, eBioscience, Cat:48-0451-82,1:200) and afterwards washed with PBS. Before sorting, the cell suspensions were stained by the live/dead marker SYTOX Blue (final concentration 1μM). Viable (SYTOX Blue negative) single cells (CD11b negative cells) were sorted by flow cytometry (SH800; Sony). Single-cells were sorted into 96 well plates filled with 4 mL lysis buffer containing 0.05% Triton X-100 (Sigma) and, ERCC (External RNA Controls Consortium) RNA spike-in Mix (Ambion,Life Technologies) (1:24000000 dilution), 2.5 mM oligo-dT, 2.5 mM dNTP and 2 U/mL of recombinant RNase inhibitor (Clontech) then spun down and frozen at −80°C. Plates were thawed and libraries prepared as described below.

### Library preparation for Smart-seq2

The 96-well plates containing the sorted single cells were first thawed and then incubated for 3 min at 72°C and thereafter immediately placed on ice. To perform reverse transcription (RT) we added each well a mix of 0.59 μL H2O, 0.5 μL SMARTScribe™ Reverse Transcriptase (Clontech), 2 μL 5x First Strand buffer, 0.25 μL Recombinant RNase Inhibitor (Clontech), 2 μL Betaine (5 M Sigma), 0.5 μL DTT (100 mM) 0.06 μL MgCl2 (1 M Sigma), 0.1 μL Template-switching oligos (TSO) (100 μM AAGCAGTGGTATCAACGCAGAGTACrGrG+G). Next RT reaction mixes were incubated at 42°C for 90 min followed by 70°C for 5 min and 10 cycles of 50°C 2 min, 42°C 2 min; finally ending with 70°C for 5 min for enzyme inactivation. Preamplification of cDNA was performed by adding 12.5 μL KAPA HiFi Hotstart 2x (KAPA Biosystems), 2.138 μL H2O, 0.25 μL ISPCR primers (10 μM, 5’ AAGCAGTGGTATCAACGCAGAGT-3), 0.1125 μL Lambda Exonuclease under the following conditions: 37°C for 30 min, 95°C for 3 min, 23 cycles of (98°C for 20 sec, 67°C for 15 sec, 72°C for 4 min), and a final extension at 72°C for 5 min. Libraries were then cleaned using AMPure bead (Beckman-Coulter) cleanup at a 0.7:1 ratio of beads to PCR product. Libraries were assessed by Bio-analyzer (Agilent 2100), using the High Sensitivity DNA analysis kit, and also fluorometrically using Qubit’s DNA HS assay kits and a Qubit 4.0 Fluorometer (Invitrogen, LifeTechnologies) to measure the concentrations. Further selection of samples was performed via qPCR assay against ubiquitin transcripts Ubb77 (primer 1 5’-GGAGAGTCCATCGTGGTTATTT-3’ primer 2 5’-ACCTCTAGGGTGATGGTCTT-3’, probe 5’-/5Cy5/TGCAGATCTTCGTGAAGACCTGAC/3IAbRQSp/-3’) measured on a LightCycler 480 Instrument II (Roche). Samples were normalized to 160 pg/μL. Sequencing libraries were constructed by using in-house produced Tn5 transposase33. Libraries were barcoded and pooled then underwent three 3 rounds of AMPure bead (Beckman-Coulter) cleanup at a 0.8:1 ratio of beads to library. Libraries were sequenced 2×150 reads base pairs (bp) paired-end on Illumina HiSeq4000 to a depth of 3×105–4×105 reads/sample.

### Processing, quality control and analyses of Smart-seq2 scRNA-seq data

BCL files were demultiplexed with the bcl2fastq software from Illumina. After quality-control with FastQC, reads were aligned using rnaSTAR^38^ to the GRCm38 (mm10) genome with ERCC synthetic RNA added. Read counts were collected using the parameter “quantMode GeneCounts” of rnaSTAR and using the unstranded values. Quantitative criteria were used to filter out low quality cells as shown in Supplementary Fig. 1b. 2538 single-cells passed the quality-control. From that point, Seurat v.3.2.3 R package was utilized^39^. Gene expressions were normalized using the SCTransform function (3000 variable features) within Seurat. The first 8 principal components (PCs) were selected based on the elbow plot and heatmap of PC embeddings, and used for downstream analysis steps. Cell type clusters were identified using the Louvain algorithm and annotated by canonical cell-type markers (Supplementary Fig. 1cd). Oligodendrocytes (2413 single cells) were extracted and analyzed separately. After processing with SCTransform (2000 variable features), the first 10 PCs were considered for downstream analyses. Unbiased clustering was performed using the Louvain algorithm that led to the identification of the 4 aforementioned oligodendrocyte populations. Gene sets of 1,2,3 and 4 were defined by using the FindMarkers function with a threshold of avg_log2FC > 1 (Fig. 1g). Gene ontology analyses were performed with the DAVID annotation tool^40^ and STRING^41^.

### Mice perfusion, cell isolation for Drop-seq

Four young (3-month old), and four old (24-month old) male C57BL/6 mice were deeply anesthetized and perfused with cold PBS. Each brain was removed, individually microdissected under a dissection microscope and dissociated in the same way as described above. To collect enough cells for loading onto the 10x Genomics Chromium chip, two gray matter/ white matter tissue samples were combined into one sample. After tissue dissociation, SYTOX Blue negative cells were sorted into a 2 mL Eppendorf tube with 1 mL RPMI+5% FBS. Sorted cells were centrifuge at 300xg for 10min at 4°C. Cell pellets were resuspended in 0.04% BSA+PBS catching media at a concentration of 700-900 cells per μl.

### Library preparation for Drop-seq

Single-cell suspensions were loaded onto the Chromium Single Cell Controller using the Chromium Single Cell 30 Library & Gel Bead Kit v3.1 (10X Genomics) chemistry following the manufacturer’s instructions. Sample processing and library preparation was performed according to manufacturer instructions using AMPure beads (Beckman Coulter). Libraries were sequenced on the DNBSEQ Sequencing System (BGI group).

### Processing and analyses of Drop-seq data

Fastq files were processed with Cell Ranger v3 and v4 for the aging and *Rag1*^-/-^ datasets, respectively, and aligned to the mm10 (Ensembl 93) genome. From that point, Seurat v.3.2.3 R package^39^ was used for the analysis of both datasets. In order to avoid batch-specific artefacts in the aging dataset, the sequenced libraries (4 sequencing runs from 8 animals) were integrated using the Seurat 3 integration workflow^42^. The first 30 PCs were selected for the downstream analyses of the integrated aging dataset. Major cell types were identified using Louvain clustering and canonical cell marker expression (Supplementary Fig.1e-f). Oligodendrocytes (6051 single cells) were extracted from the dataset and integrated by batch separately. Before the integration, each batch of oligodendrocytes were normalized with SCTransform. For the integrated dataset, the first 10 PCs were considered for further analyses steps. Unbiased clustering using the Louvain algorithm identified the aforementioned 7 oligodendrocyte populations. The *Rag1*^-/-^ dataset was normalized with the SCTransform function and further processed as described for the aging dataset; identifying the major cell types by matching canonical cell type markers with cluster specific marker genes. Oligodendrocytes were analysed separately; the first 10 PCs were selected following normalization with SCTransform. Unbiased clustering identified the same clusters in the aging dataset, with the exception of IRO population. All differential gene expression analyses were conducted using the FindMarkers function with the Wilcoxon Rank Sum test for both 10X datasets.

### Immunohistochemistry

Animals were anesthetized by 10 mg/ml ketamine and 1 mg/ml xylazine solution i.p. perfused transcardially with 4% paraformaldehyde (PFA). Post fixation of brain tissue was done in 4% PFA overnight. Then the brain tissue was further cryo-protected in 30% sucrose in PBS for 24 h. After freezing the tissue on dry ice using Tissue-Tek O.C.T, 30 μm coronal sections were cut by cryostat Leica CM 1900. Free-floating sections were collected in a solution containing 25% glycerol and 25% ethylenglycol in PBS. The sections were rinsed with 1x PBS containing 0.2% Tween-20 and permeabilized in 0.5% Triton X-100 for 30 min. Fab fragment goat anti mouse IgG (1:100) (Dianova) was added for 1 h at room temperature to block endogenous mouse tissue immunoglobulins. After a brief wash the sections were blocked for 1 h at room temperature in a solution containing 2.5% FCS, 2.5% BSA and 2.5% fish gelatin in PBS.

Primary antibodies, diluted in 10% blocking solution, were incubated overnight at 4°C. On the following day sections were incubated with secondary antibodies, diluted in 10% blocking solution, for 2 h at room temperature. The sections were washed with PBS followed DAPI incubation in 1x PBS for 10 minutes and mounted. The following antibodies were used: The following antibodies were used: mouse anti APC (Millipore OP80-100UG, 1:100), rabbit anti B2m (abcam ab75853-100ul, 1:100), rabbit anti STAT1 (cell signalling 14994S,1:250), Rat anti CD8 (promega 100702,1:100) rabbit anti Iba1 (wako 234 004, 1:250) goat anti Serpina3n (bio-techne AF4709, 1:100) rat anti C4b (thermo fisher scientific MA1-40047,1:25) rabbit anti Olig2 (millipore AB9610, 1:250) mouse anti-Gstp (BD 610719, 1:250), anti mouse 555 (thermo fisher scientific A-21422, 1;500) anti mouse 647 (thermo fisher scientific A-21235, 1:500) anti mouse 488 thermo fisher scientific A-21202, 1:500) anti rabbit 555 (thermo fisher scientific A-21428, 1:500) anti rabbit 488 (thermo fisher scientific A-11008,1:500) anti rat 555 (thermo fisher scientific A-21434, 1:5600) anti goat 555 (thermo fisher scientific A-32116, 1:500). For CC1, B2m, Gstp, OLIG2, STAT1, Serpina3n staining, antigen retrieval protocol using citrate buffer (10 mM, pH 6) was performed on free-floating sections followed by staining protocol as mentioned above. For CD8e, STAT1 and CC1 combined staining, the sections were permeabilized with 0.5% Triton X-100 for 30 min at room temperature and blocked in 5% goat serum (Vector Laboratories, 1:200), goat anti-rat 488 and goat anti-mouse 647 (Invitrogen, 1:500) for 1 h at room temperature. Sections were then washed with PBS and incubated with streptavidin 555 (Invitrogen, 1:500). To determine the proximity of CD8^+^ T cells to STAT1^+^CC1^+^ Oligodendrocytes (IRO), CD8+ T cells (or random DAPI^+^ cell) were selected manually, from which a 20 μm circle was drawn from the center of the cell via Image J automatically. We then quantified the percentage of STAT1^+^CC1^+^ oligodendrocyte located within that circle. Conversely, we proceeded similarly but took oligodendrocytes as a reference and quantified the T cells within that 20 μm perimeter. Images were acquired via a Leica TCS SP5 confocal microscope or with LSM900 Zeiss microscope and were processed and analyzed with ImageJ 1.41 image processing software.

### Sub-regional localization analysis

To compare the difference between frontal white matter and medial white matter, coronal sections with corpus callosum were divided into two groups following the Allen Mouse Brain Atlas: the white matter in the front of the brain which does not contain lateral ventricles was defined to be “frontal white matter”, and white matter from sections which have lateral ventricles is defined to be “medial white matter”. For each quantification, two brain sections were selected from each group of 3-4 mice. For statistical analysis paired t test was used.

### RNAscope in Situ Hybridization

RNAscope in situ hybridization assay was applied to detect mRNA of MBP in the brain cryosections prepared from aged wild-type mice was done as performed previously^16^. Briefly, the assay was performed using a commercially available kit, RNAscope Multiplex Fluorescent Detection Reagents v2 (Advanced Cell Diagnostics, ACD) and the manufacturer instruction was followed. Briefly, 30 μm cryosections were fixed on superfrost plus slides; they were pretreated with hydrogenproxide for 10 min at room temperature and then with antigen retrieval reagent (5min boiling) to unmask the target RNA. After applying Protease III on the sections for 30 min at 40°C, probe hybridization was done by incubating sections in mouse MBP probe assigned to channel 3, diluted 1:50 in probe diluent, for 2 h at 40°C. Afterward, signal amplification and detection were performed according to the instruction of the kit. Signal detection was done using Opal dyes (Opal520-green) diluted 1:3000 in TSA buffer. To visualize microglia, after in situ hybridization, immunohistochemistry assay was performed using Iba1 antibody (Wako, 1:1000). After washing with 1x PBS sections were incubated for 30 second with 1xTrueBlack to remove autofluorescence background. The nuclei of cells were counterstained with DAPI (4’,6-diamidino-2-phenylindole) and mounted. Images were acquired via a Leica TCS SP5 confocal microscope or with LSM900 Zeiss microscope and were processed and analyzed with ImageJ 1.41 image processing software.

### Statistical analysis

For immunohistochemistry analysis 3-6 sections from each animal were analyzed. Data are shown as mean ± error of the mean (sem). No statistical methods were used to predetermine sample sizes, but our sample sizes are similar to those generally employed in the field. Each dot represents one animal. Normal distribution of the samples was tested by Shapiro-Wilk test. For statistical analysis paired or unpaired T test, or Mann-Whitney were used to compare two groups. One-way ANOVA followed by uncorrected Fisher’s LSD or Dunnett’s multiple comparisons test or Kruskal-Wallis followed uncorrected Dunn’s test were used for multiple comparison. Test were chosen accordingly to their distribution. In all tests a p value of < 0.05 was considered as significant. Statistical analyses were done using GraphPad Prism (GraphPad Software, Inc.).

### Data availability

The scRNA-seq data will also be deposited at GEO (NCBI) with the revised version. All other data that supports findings are available upon request from the authors.

### Code availability

The R code used for the analyses can be provided upon request from the authors.

